# Comprehensive analysis of potential immunotherapy genomic biomarkers in 1,000 Chinese patients with cancer

**DOI:** 10.1101/366062

**Authors:** Shunchang Jiao, Yuansheng Zang, Chun Dai, Xiaoman Xu, Xin Cai, Guan Wang, Jinwang Wei, Angela Wu, Wending Sun, Qiang Xu

**Author notes:** Shunchang Jiao and Yuansheng Zang contribute equally to this work. Correspondence: Dr. Qiang Xu, GenomiCare Biotechnology (Shanghai) Co., Ltd., 5th Floor, Building #2, No. 111 Xiangke Road, Shanghai, China 201210.

## Abstract

**Background:** Tumor mutation burden (TMB), DNA mismatch repair deficiency (dMMR), microsatellite instability (MSI), and PD-L1 amplification (PD-L1 AMP) may predict the efficacy of PD-1/PD-L1 blockade. In this study, we aimed to characterize the distributions of these biomarkers in over 1,000 Chinese patients with cancer.

**Methods:** TMB, MSI, dMMR, and PD-L1 AMP were determined based on whole-exome sequencing of tumor/blood samples from > 1,000 Chinese patients with cancer.

**Results:** Incidence rates among 953 Chinese patients with cancer showing high TMB (TMB-H), high MSI (MSI-H), dMMR and PD-L1 AMP were 35%, 4%, 0.53% and 3.79%, respectively. We found higher rates of TMB-H among hepatocellular carcinoma, breast cancer, and esophageal cancer patients than was reported for The Cancer Genome Atlas data. Lung cancer patients with *EGFR* mutations had significantly lower TMB values than those with wild-type *EGFR*, and increased TMB was significantly associated with dMMR in colorectal cancer (CRC). The frequency of tumors with MSI-H was highest in CRC (14%) and gastric cancer (4%). PD-L1 AMP occurred most frequently in lung squamous cell carcinoma (14.3%) and HER2-positive breast cancer (8.8%). Most MSI-H and dMMR cases exhibited TMB-H, but the overlap among the other biomarkers was low.

**Conclusion:** While MSI and dMMR are associated with higher mutational loads, correlations between TMB-H and other biomarkers, between MSI-H and dMMR, and between PD-L1 AMP and other biomarkers were low, indicating different underlying causes of the four biomarkers. Thus, it is recommended that all four biomarkers be assessed for certain cancers before administration of PD-1/PD-L1 blockade treatment.

## Introduction

The PD-1/PD-L1 blockade has become a powerful approach for treating multiple types of cancer. Many patients benefit from such treatments, exhibiting not only a higher objective response rate but also durable remission for many years.^1-7^ While treatment with PD-1/PD-L1 blockade can be highly effective, not every patient or cancer type responds to these inhibitors ^8^, and some patients experience hyper progression after immunotherapy.^9^ Identifying which patients may benefit from these inhibitors is one of the most significant current challenges. Therefore, the identification of precise biomarkers to predict the efficacy of PD-1/PD-L1 blockade is critical.

Recently, five clinical trials across 15 cancer types involving 149 patients with high microsatellite instability (MSI-H) or DNA mismatch repair deficiency (dMMR) reported complete or partial response to pembrolizumab in 39.6% of patients. Moreover, among 78% of the responding patients, the response lasted for 6 months or longer.^10, 11^ These results accelerated the FDA approval of this treatment for unresectable or metastatic solid tumors in patients with MSI-H or dMMR.^12^

Another emerging biomarker is high tumor mutation burden (TMB-H), which is closely related to the PD-1/PD-L1 blockade response. Melanomas have the highest mutational loads among human tumors and also exhibit an unusually high response rate of 30–40% to the PD-1/PD-L1 blockade. Urothelial carcinoma, NSCLC, and head and neck squamous cell carcinoma have the next highest median mutational loads and respond to the PD-1/PD-L1 blockade at rates above 15%. In contrast, glioblastoma and pancreatic, ovarian, and prostate cancer, which have relatively low median mutational loads, show little response to PD-1 blockade treatment.^13^

Finally, research has identified PD-L1 (CD274) amplification (PD-L1 AMP) as a predictor of response to PD-1/PD-L1 blockade therapy. In classical Hodgkin lymphoma, 97% of patients exhibit PD-L1 AMP.^14^ Compared with other cancers, patients with Hodgkin lymphoma exhibit a higher overall response rate, reaching 69%.^15, 16^ In rare metastatic basal cell carcinoma, PD-L1 AMP is also related to response to nivolumab.^17^ Thus, there is a strong correlation between PD-L1 AMP and response to the PD-1/PD-L1 blockade.

Given that the above markers are potential biomarkers of PD-1/PD-L1 blockade efficacy, we aimed to characterize the distributions of these biomarkers in over 1,000 Chinese patients with cancer using exome profiling data. We also explored the relationships among TMB, MSI-H, dMMR, and PD-L1 AMP.

## Methods

### Patient characteristics

We collected over 1,000 Chinese cancer specimens from more than 70 hospitals in 20 Chinese provinces from October 2015 to March 2016. In total, there were 1,179 samples, including 524 biopsy samples and 655 formalin-fixed, paraffin-embedded (FFPE) samples. Blood samples were also collected as controls. All procedures followed the Molecular Pathology Clinical Practice Guidelines and Reports^18^ and were performed in accordance with the 1964 Declaration of Helsinki. Informed consent was obtained from each of the participants.

### Whole-exome sequencing (WES) analysis

The complete exomes of tumor samples and matched blood samples were sequenced in each patient. DNA was fragmented and hybridized to the SureSelect Human All Exome Kit V5 (Agilent Technologies, Santa Clara, CA, USA), containing exon sequences from 27,000 genes. Exome shotgun libraries were sequenced on the Illumina Xten platform, generating paired-end reads of 150 bp at each end. Image analysis and base calling were performed with CAVSAVR (Illumina, San Diego, CA, USA) using default parameters. Sequencing adaptors and low-quality reads were removed to obtain high-quality reads. These were aligned to the NCBI human reference genome hg19 using the Burrows-Wheeler Aligner alignment algorithm.^19^

We used the Genome Analysis Toolkit (GATK, version 3.5) to process reads. Localized insertion-deletion (indel) realignments were performed by GATK. GATK Realigner Target Creator was used to identify regions for realignment. For single-nucleotide variant (SNV) calling, the MuTect algorithm was applied to identify candidate somatic SNVs in tumors by comparison with the matched control blood sample from each patient. SNV annotation was performed by ANNOVAR. We used dbNSFP31 to predict nonsynonymous mutations in the encoded proteins. For dMMR, we identified SNVs in *MLH1*, *MSH2*, *MSH6,* and *PMS2*.

For indel detection, tumor and blood samples were analyzed with VarscanIndel. Candidate somatic indels were identified based on the following criteria: (1) supported by at least five reads and (2) the number of supporting reads divided by the maximum read depth at the left, and right breakpoint positions were > 0.05. All somatic indel calls were manually checked with the Integrative Genomics Viewer.

The CNVnator software tool was used to detect somatic copy number variations (CNVs).^20^ All parameters were set to their defaults for filtering samples, and the bin size was set to 50–60 according to the average coverage depth. A PD-L1 copy number ≥ 3 was defined as PD-L1 AMP.

### TMB evaluation

TMB was defined by the total number of nonsynonymous somatic mutations (NSM), which was determined by comparing sequence data between tumor tissues and matched blood samples using a previously described method.^21^ We defined the higher tertile of the TMB of each cancer type as the threshold for TMB-H according to the method of a prior study.^22^

### MSI evaluation

All autosomal microsatellite tracts containing five or more repeating subunits 1–5 bp in length in GRCh37/hg19 were identified using MISA (http://pgrc.ipk-gatersleben.de/misa/misa.html). Detailed calculations were performed as described previously.^23^ Patients were classified into the microsatellite stable (MSS), low MSI (MSI-L), and high MSI (MSI-H) groups with 0–1%, 1–3.5%, and ≥3.5% unstable microsatellite sites, respectively, according to a previous method.^24^

## Results

### Cohort description

We collected 1,197 tumor samples from Chinese patients. Due to unqualified samples and sequencing failures, 953 samples were successfully sequenced by WES. These tumors encompassed four principal tumor types, including CRC and lung, breast, and gastric cancer, accounting for up to 78% of the tumors. There were also more than eight other types of tumor (Figure 1). Based on the cancer distribution in China,^25^ we collected more lung adenocarcinoma (LUAD) cases (n = 172) than lung squamous cell carcinoma (LUSC) cases (n = 42) and other subtypes (n = 37). Among breast cancer cases, the number of patients with HER2^+^ (n = 34), HER2^−^ (n = 49), and HER2 status unknown (n = 68) cancer were similar.

**Figure 1.**
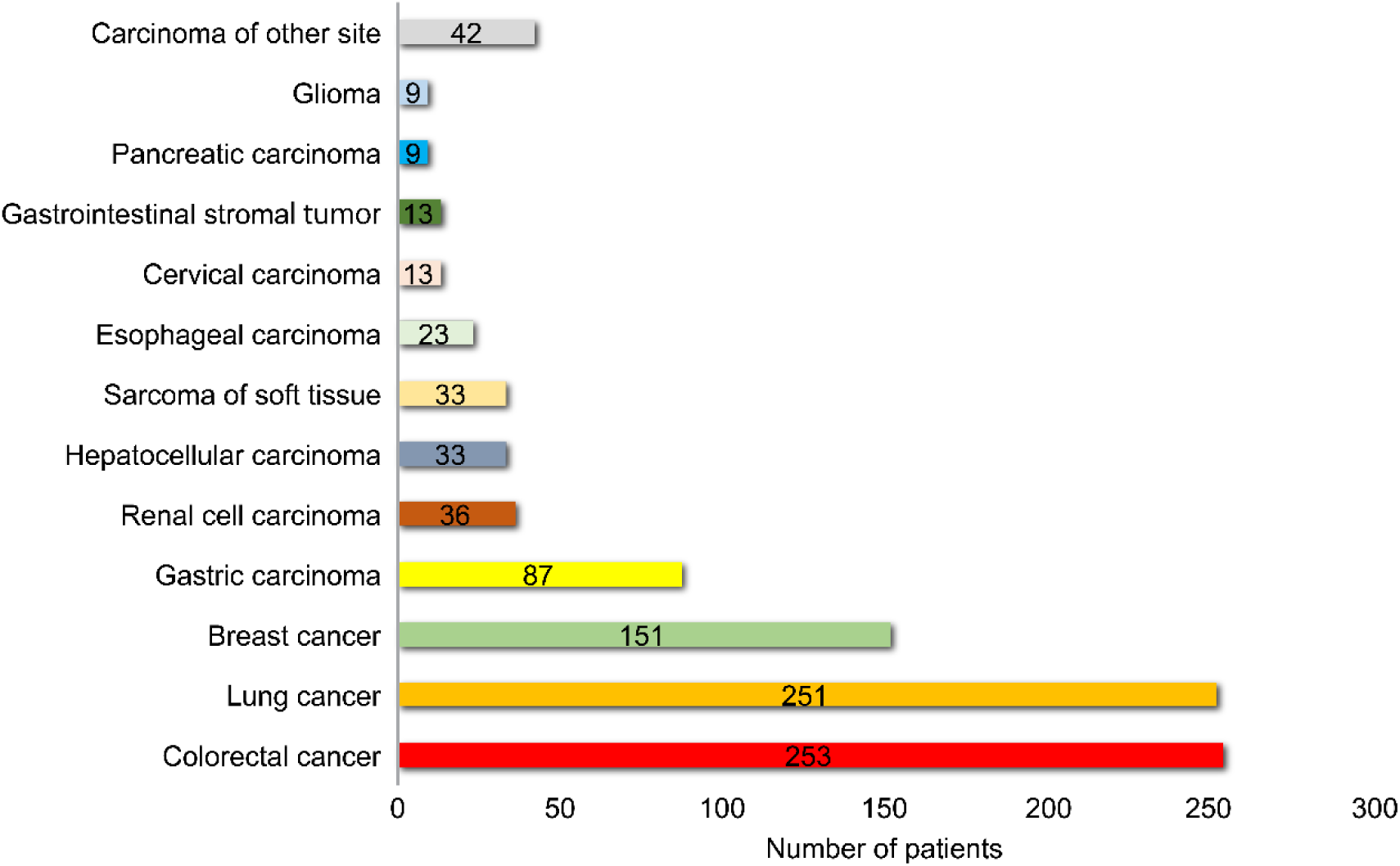
Distribution of cancer types among 953 Chinese patients analyzed by WES.

### TMB profiling

We analyzed the TMB of the 953 Chinese patients with cancer. The median TMB of each cancer type ranged from 36 to 273 NSM, with an overall median of 95 NSM, and 54 patients exhibited NSM values > 1000. The two cancer types with the highest TMB were lung cancer and CRC, with median TMB of 176 and 108 NSM, respectively. Hepatocellular carcinoma, renal cell carcinoma, and glioma also had high TMB values, while the lowest TMB was observed among gastrointestinal stromal tumors. We also found that different tumor subtypes exhibited different TMB, especially in lung cancer: LUSC cases had median TMB values more than three times higher than those in LUAD cases (273 vs. 74 NSM, respectively) (Figure 2A).

**Figure 2.**
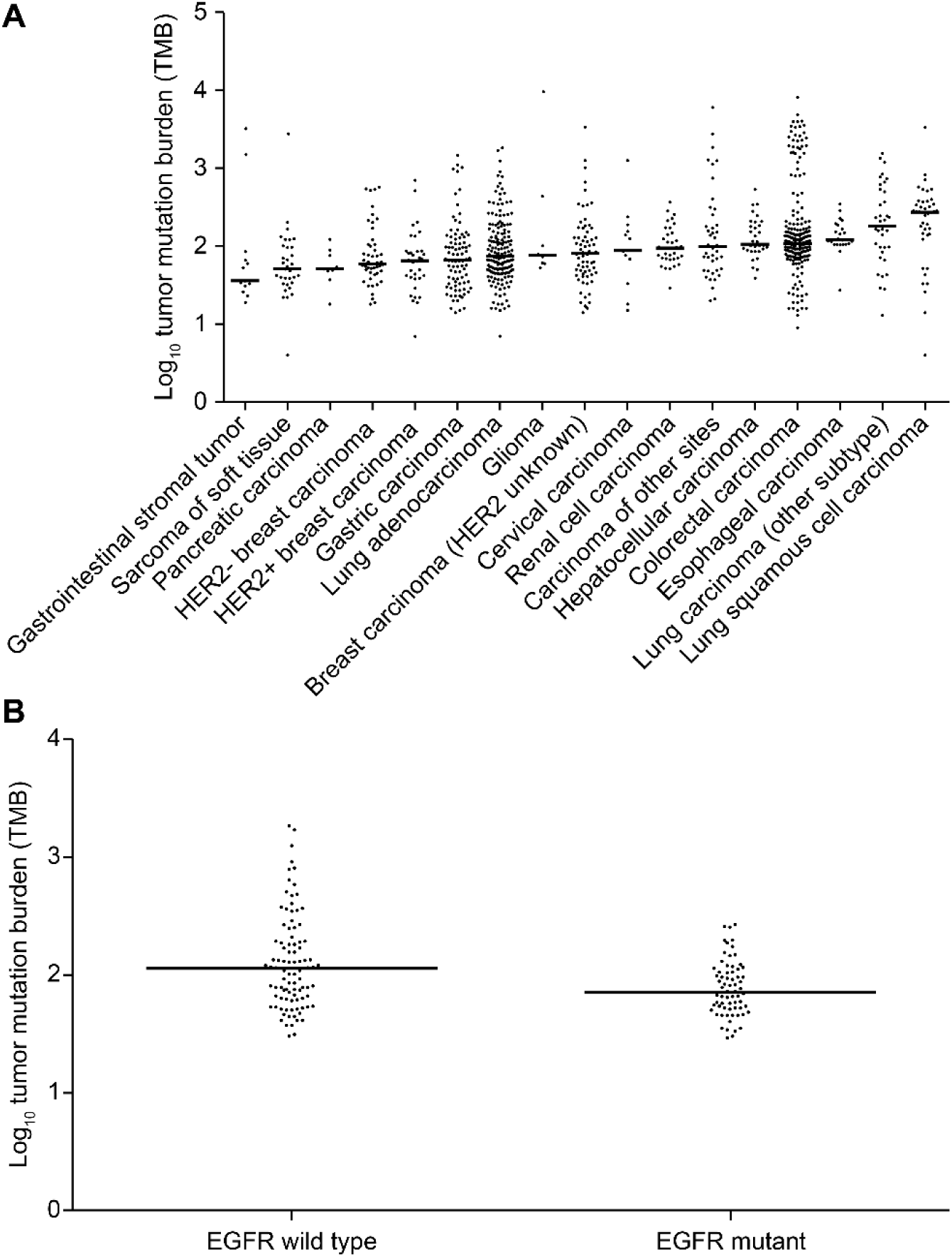
Characterization of tumor mutation burden (TMB). A. The median number of somatic nonsynonymous mutations (log_10_-transformed) was plotted for each cancer type after analysis of 953 Chinese patients. The difference in median TMB between LUAD and LUSC was statistically significant (*P* < 0.0001 by Mann-Whitney *U* test). B. The median number of somatic nonsynonymous mutations (log_10_-tranformed) was plotted for LUAD patients with or without *EGFR* mutation (*P* = 0.0039 by Mann-Whitney *U* test).

Although PD-1/PD-L1 blockade has been approved in NSCLC, some clinical trials have reported that NSCLC patients with *EGFR* mutations do not benefit from this therapy and that it may even lead to a more rapid disease progression.^2, 10, 26^ Therefore, we compared the median TMB of LUAD patients with mutant (n = 77) and wild-type (n = 99) *EGFR* genes. Patients with *EGFR* mutations had significantly lower median TMB compared with those with wild-type *EGFR* (74 vs. 113 NSM, respectively; *P* = 0.0039) (Figure 2B).

### MSI and dMMR distributions

We investigated the MSI and dMMR distributions in Chinese patients with cancer. Tumors were classified into three groups based on the proportion of unstable microsatellites, consistent with previous reports^24^: MSS, below 0–1% (n = 597); MSI-L, 1–3.5% (n = 12); and MSI-H, ≥3.5% (n = 28). The frequency of tumors classified as MSI-H was 14% for CRC and 4% for gastric cancer, while lung and breast cancer exhibited MSI-H frequencies below 3% (Figure 3A). Gastrointestinal stromal tumors exhibited a high frequency of dMMR, while those of hepatocellular carcinoma, sarcoma of soft tissue, and renal cell carcinoma were among the lowest. The dMMR frequency of lung carcinoma (other types) was nearly double those in LUAD and LUSC. HER2^−^ breast carcinoma exhibited a higher dMMR frequency than HER2^+^/ HER2 unknown breast carcinoma (Figure 3B).

**Figure 3.**
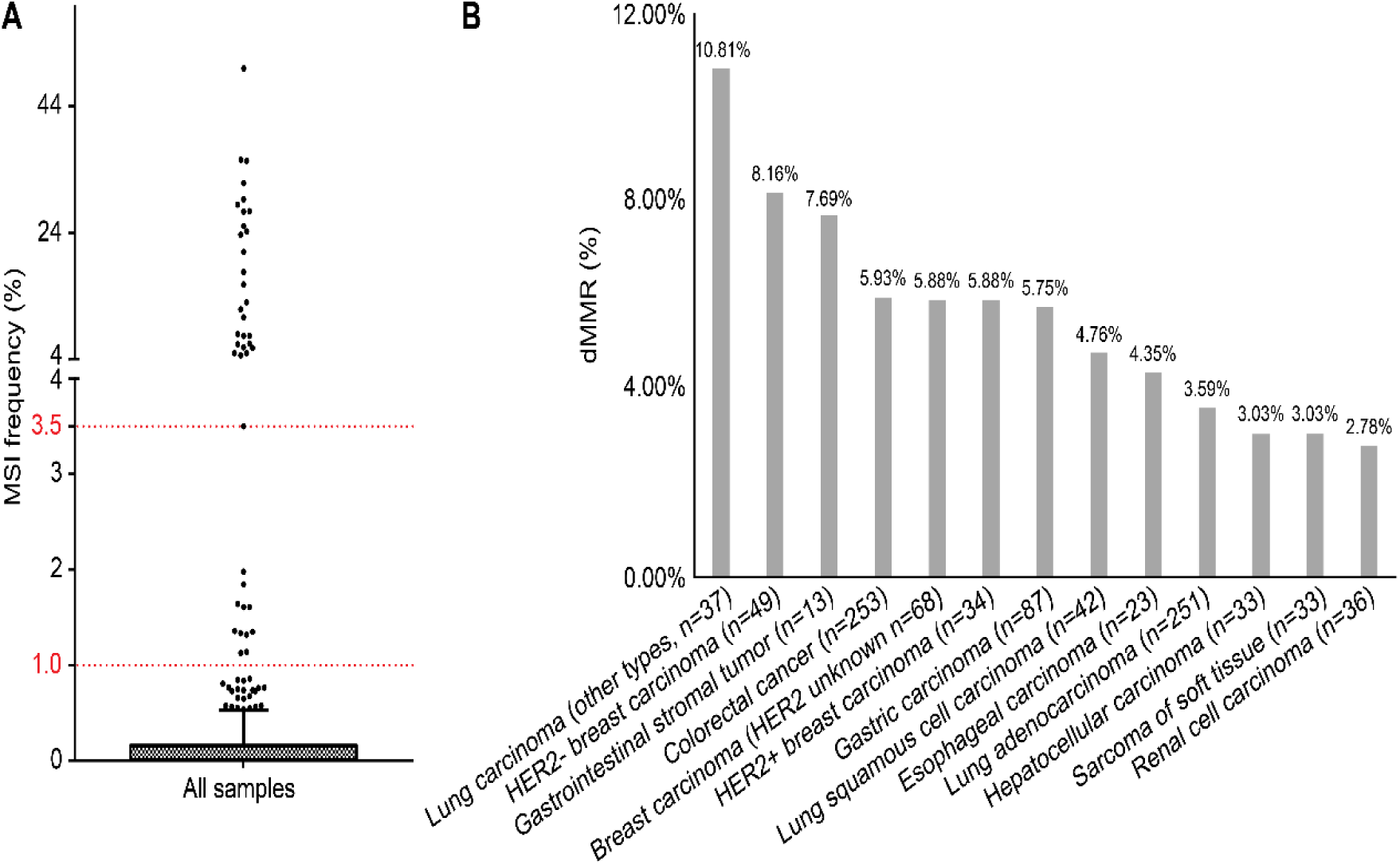
MSI and dMMR distribution in 953 Chinese patients. A. MSI distribution. Patients in the MSS, MSI-L, and MSI-H groups exhibited 0–1%, 1–3.5%, and ≥3.5% unstable microsatellite sites, respectively. B. dMMR distribution.

### PD-L1 AMP distribution

We analyzed the distribution of PD-L1 AMP among Chinese patients with cancer. We observed that PD-L1 AMP occurred most frequently in LUSC (14.3%), and HER2^+^ breast cancer (8.8%), and breast cancer with unknown HER2 status (5.8%). In contrast, LUAD and CRC had lower rates of PD-L1 AMP at 1.75% and 1.59%, respectively (Figure 4). Thus, LUAD and LUSC exhibit large differences in levels of PD-L1 AMP.

**Figure 4.**
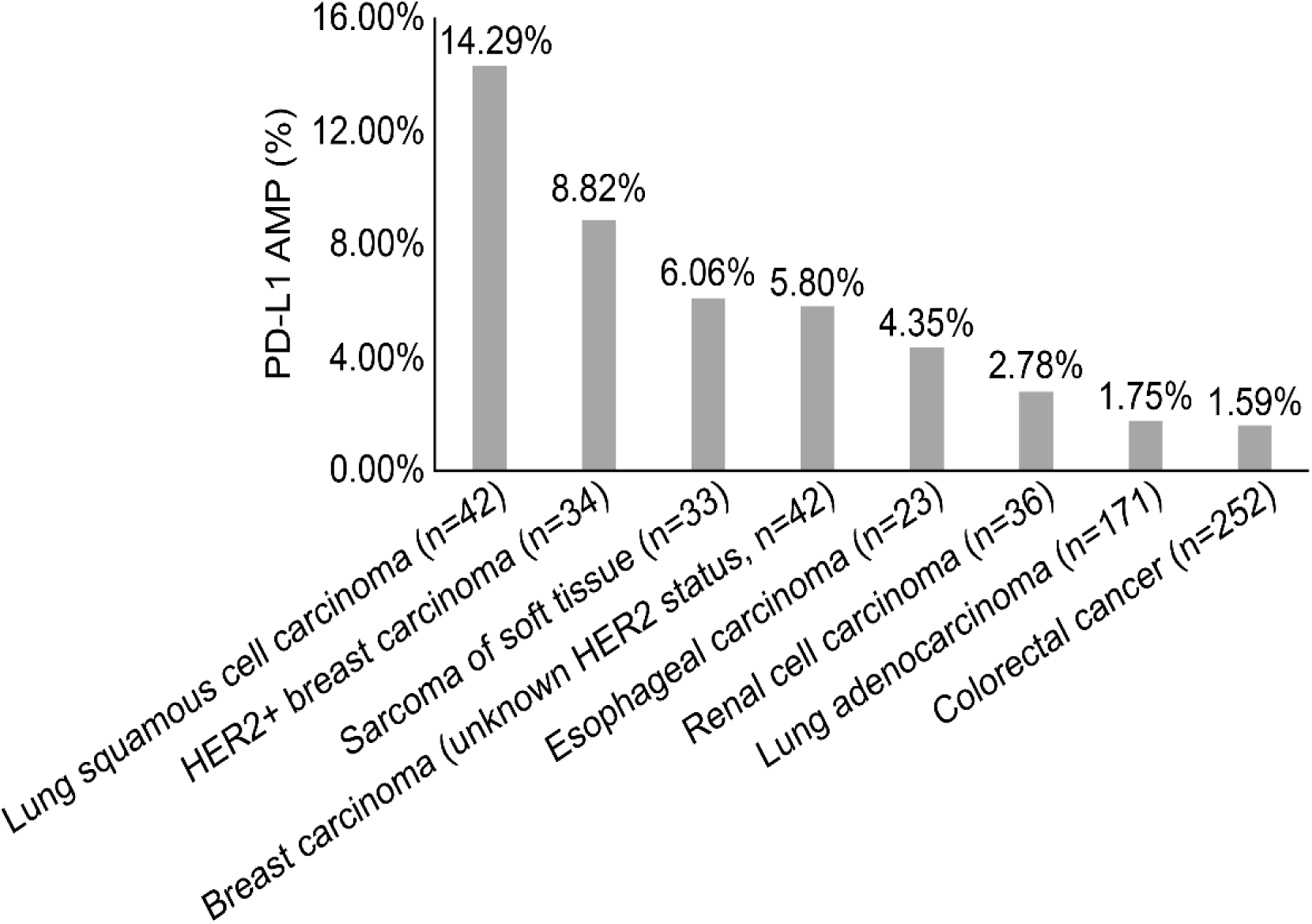
PD-L1 amplification frequency across eight major tumor types.

### Relationships among biomarkers

In addition to exploring the distributions of the four biomarkers in Chinese patients with a wide variety of tumor types, we also analyzed the correlations among these biomarkers. As shown in Figure 5A, the four biomarkers overlap with each other. Among 337 Chinese patients with cancer showing TMB-H, 11.28% are also positive for dMMR, 7.4% for MSI-H, and 2.7% for PD-L1 AMP. In addition, 9 cases in the TMB-H cohort also exhibited MSI-H and dMMR simultaneously, while one patient was positive for all four biomarkers. Up to 76.26% of TMB-H tumors did not coincide with any of the other biomarkers (Figure 5B). In conclusion, TMB-H is a relatively independent biomarker with a small overlap with other biomarkers.

**Figure 5.**
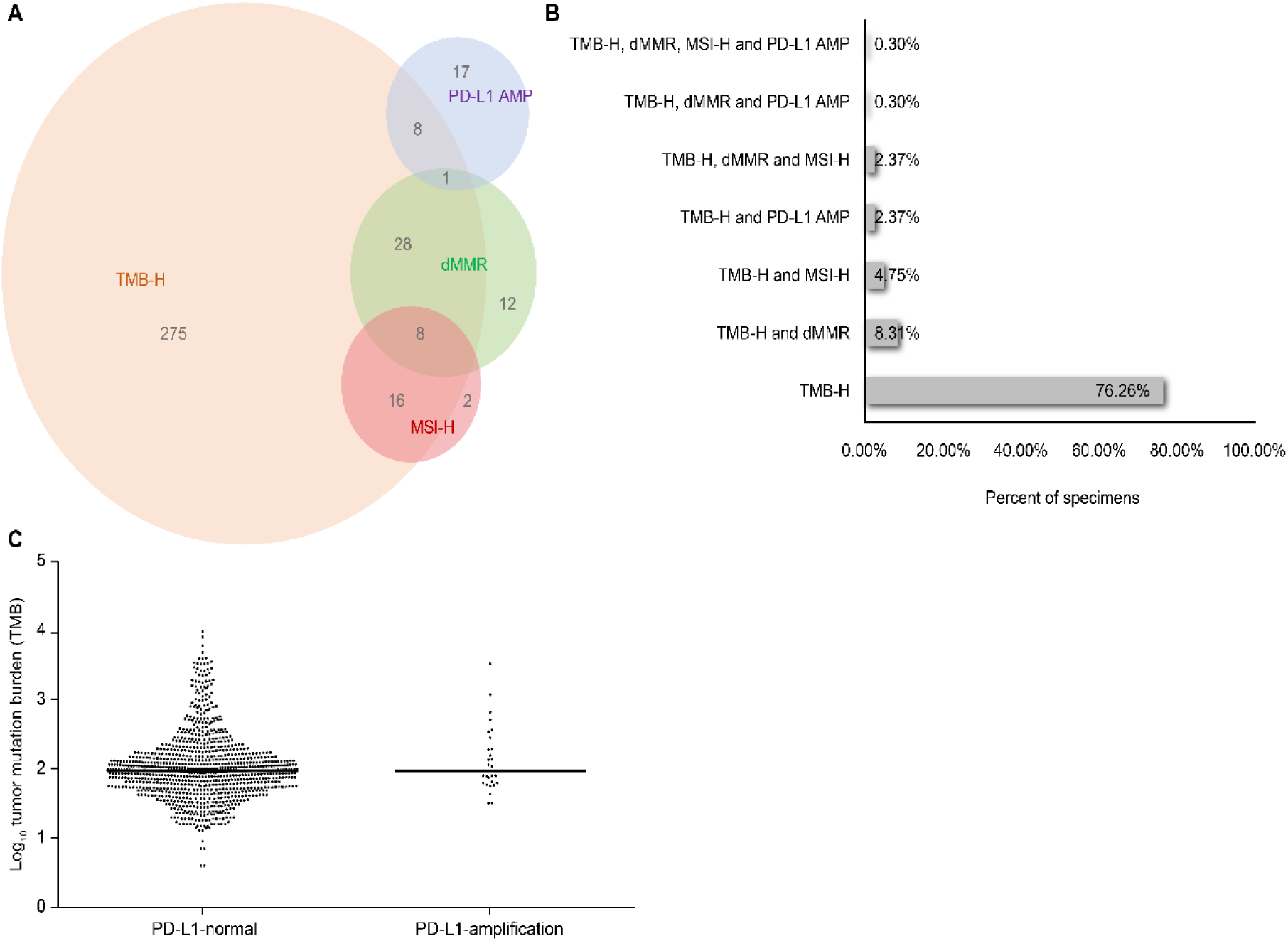
Relationship between TMB-H and MSI, dMMR, and PD-L1 AMP across cancer types in Chinese patients. A. Overlapping distributions of TMB, MSI, dMMR and PD-L1 AMP. B. Proportions of TMB-H patients positive for other biomarkers. C. TMB in all samples with (n = 28) or without (n = 922) CD274 (PD-L1) amplification.

We also calculated the TMB values of patients with amplified and normal PD-L1. The median values of TMB in the two groups were 94 and 95 NSM, respectively, reflecting no significant difference between the two groups (Figure 5C). Moreover, 34.6% of PD-L1 AMP patients were classified as TMB-H. These results suggest that the correlation between TMB-H and PD-L1 AMP is small. Thus, it is necessary to simultaneously detect both in order to identify more suitable cancer patients for immune therapy.

Furthermore, we investigated the correlations between MSI-H and the other three biomarkers. MSI arises from mutations or epigenetic alterations in the MMR proteins (MLH1, MSH2, MSH3, MSH6, PMS1, or PMS2).^27, 28^ As anticipated, we found a correlation between MSI-H and dMMR: 32.14% of Chinese patients with cancer showing MSI-H exhibited dMMR, and those cancer patients were also TMB-H (Figure 6A). Thus, while MSI-H and dMMR were correlated, the association was not particularly strong, suggesting that MSI-H may also be caused by additional factors.

**Figure 6.**
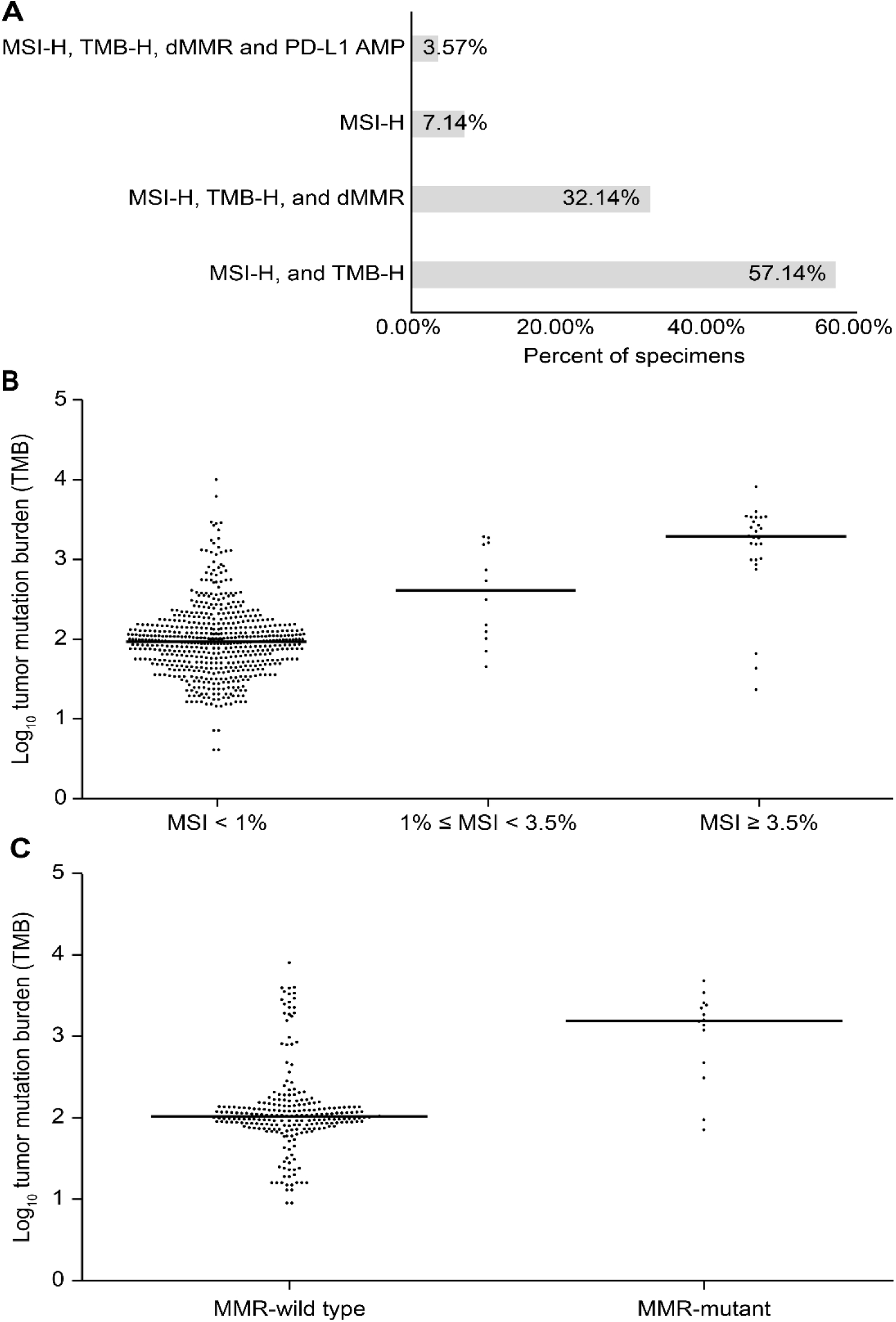
Correlations between MSI-H and TMB-H, dMMR, and PD-L1 amplification. A. Proportions of MSI-H cases with other biomarkers. B. TMB in the MSI-H, MSI-L, and MSS groups. Significant *P*-values were observed by Mann-Whitney *U* test between MSI-H and MSI-L (*P* = 0.0062), MSI-H and MSS (*P* < 0.001), and MSI-L and MSS (*P* = 0.0058). C. TMB in CRC with (n = 15) or without (n = 237) MMR mutations. *P* < 0.0001 by Mann-Whitney *U* test.

MSI-H tumors exhibit high levels of neoantigen, causing strong local and systemic immune responses.^29^ Therefore, it is necessary to investigate the relationship between MSI and TMB. The median TMB values of the MSI-H, MSI-L, and MSS groups were 1945, 408, and 90 NSM per tumor, respectively, reflecting significant differences among groups (Figure 6B). About 89% of Chinese patients with cancer exhibiting MSI-H status were classified in the TMB-H group.

We also found a remarkable correlation between dMMR and TMB-H in CRC A comparison of the TMB values of dMMR patients (n = 15) and MMR-proficient patients (n = 237) showed that increased TMB was significantly associated with dMMR (*P* < 0.0001). dMMR tumors on average had 15-fold more NSM (1567 vs. 106 NSM) than MMR-proficient patients (Figure 6C).

In each of the MSI-H and dMMR groups, there were only two cases with PD-L1 AMP, representing rates of 7% and 4%, respectively. PD-L1 AMP cases also exhibited low frequencies of dMMR and MSI-H (<10% each), which is consistent with a previous report showing that among 365 solid tumors only the 10.9% of MSI-H cases exhibited PD-L1 AMP.^30^

## Discussion

To our knowledge, this is the first study characterizing TMB, MSI, dMMR, and PD-L1 AMP in Chinese patients with more than 18 tumor types, as determined by WES. The incidence rates in TMB-H, MSI-H, dMMR, and PD-L1 AMP were 35%, 4%, 0.53% and 3.79 % respectively.

Chinese patients with cancer exhibited a characteristic pattern of TMB. The two cancer types with the highest TMB were lung cancer and CRC, similar to the results of prior studies.^31, 32^ In contrast, hepatocellular carcinoma, breast cancer, and esophageal cancer cases exhibited TMB values 1–3 times higher than those in The Cancer Genome Atlas (TCGA) cohort.^33^ The CHECKMATE-040 and APHINITY trials showed that adjuvant treatment with nivolumab and pertuzumab effectively improved the overall response rate of hepatocellular carcinoma previously treated with sorafenib and the invasive disease-free survival of HER2^+^ breast cancer.^34, 35^ This indicates that more Chinese hepatocellular carcinoma, breast cancer, and esophageal cancer patients could benefit from the PD-1/PD-L1 blockade and that TMB can be used as a potential immunotherapeutic biomarker.

We collected more LUAD patients than LUSC patients, and we found that the TMB values of two types differed among Chinese patients. In contrast, the prevalence of TMB-H among LUAD patients is similar to that among LUSC patients in TCGA.^33^ Moreover, in clinical trials, the treatment of both advanced squamous and non-squamous NSCLC with nivolumab resulted in similar improvements over treatment with docetaxel, with overall survival rates of 42% and 51% and response rates of 20% and 19%, respectively.^2, 36^ Similarly, in a trial with nivolumab as the first-line treatment for advanced NSCLC, squamous and non-squamous groups had similar median progression-free and overall survival.^37^ Based on our findings, however, Chinese patients with LUSC may demonstrate the higher potential for clinical benefits of the PD-1/PD-L1 blockade. In addition, we found that patients with *EGFR* mutations had significantly lower median TMB compared with those with wild-type *EGFR*, consistent with another study in which the TMB value of *EGFR*-mutant non-squamous NSCLC was half that of wild-type.^38^ This result may help to explain why in *EGFR*-mutant advanced NSCLC, immune checkpoint blockade does not improve overall survival over that achieved with docetaxel alone.

Several studies have analyzed the correlation between TMB and the efficacy of immunotherapy in detail, and all have shown high concordance between TMB and treatment efficacy.^13, 39, 40^ A study containing 151 cancer patients further confirmed a linear relationship between TMB and the clinical outcome of immunotherapy.^39^ Moreover, the cut-off value of TMB to predict treatment response has been determined in several studies. Another study involving melanoma and NSCLC clinical data found the cut-off to be 192 NSM, with a 74% sensitivity and 59.3% specificity.^31^ In this study, more than 20% of patients with LUSC, other subtypes of lung carcinoma, hepatocellular carcinoma, glioma, esophageal carcinoma, and CRC had NSM counts above 192, indicating that TMB-H may be a good indicator for the use of PD-1/PD-L1 blockade. However, the optimal TMB cut-offs require further exploration in clinical trials with Chinese patients with cancer in order to guide the use of immunotherapy.

In the current study, the frequency of tumors classified as MSI-H was 14% for CRC, while lung and breast cancer exhibited MSI-H frequencies below 3%. By contrast, a comprehensive review of studies that investigated MSI before 2014 showed that CRC exhibited MSI-H frequencies >10%, while that of lung cancer was <2%.^41^ Although the distributions of MSI-H in Chinese patients with cancer observed in this study were consistent with those in cancer patients in other countries, the frequency of MSI-H in gastric cancer is very low (4%), and the rates of MSI-H in cancer patients from other countries were 18–22%.^41, 42^ This suggests that MSI-H may not be useful in screening gastric patients before PD-1/PD-L1blocker and that other biomarkers or a combination of MSI-H with other biomarkers should be investigated further.

Compared with TMB and MSI, research involving the frequency of PD-L1 AMP in many cancer types is lacking. In this study, we fully surveyed PD-L1 AMP in Chinese patients with cancer. We found relatively high rates in LUSC (14.3%), HER2^+^ breast cancer (8.8%), and breast cancer with unknown HER2 status (5.8%), and low rates in LUAD (1.75%) and CRC (1.59%). In other studies, rates of PD-L1 AMP of 29% in triple-negative breast cancer, 5% in glioblastoma, and 3% in CRC have been observed.^43, 44^ A study of more than 100,000 patients with many cancer types also surveyed the PD-L1 AMP distribution.^45^ In this study, sarcomatoid kidney carcinoma and nasopharyngeal cancer exhibited rates of PD-L1 AMP over 5%, while LUAD, CRC, and breast cancer exhibited rates of 0.6%, 0.2%, and 2%, respectively. Thus, rates of PD-L1 AMP among Chinese patients with LUAD, breast cancer, and CRC were similar to those observed in other studies.

While TMB, dMMR, MSI, and PD-L1 AMP are different genetic alterations that occur in many cancers, they may be inherently related. In an analysis of 11,348 cancer patients, 27% of patients with TMB-H exhibited MSI-H, and 70% of MSI-H cases had high TMB.^46^ An analysis of 100,000 cancer genomes from TCGA, found that the 83% of MSI-H samples had high TMB.^21^ By contrast, in this study, we found that Chinese cancer patients showing TMB-H exhibited relatively low rates of MSI-H and dMMR. On the other hand, about the 89% of Chinese cancer patients with an MSI-H status belonged to the TMB-H group. These results indicate that MSI-H can cause TMB-H, but TMB-H is not primarily caused by MSI-H.

We found that 32.14% of Chinese patients with cancer showing MSI-H exhibited dMMR. This overlap between the MSI-H and dMMR cases is higher than what was previously reported (13.37%).^42^ We also found that, on average, dMMR CRC carried 15 folds more NSM (1567 vs. 106 NSM) than MMR-proficient CRC, in accordance with a previous report (1782 vs. 73 NSM, respectively).^11^

These findings can be interpreted based on data from a previous study regarding the hypermutation of human cancers. This study found that TMB of >100 Mut/Mb may be classified as MSS but contain MMR and replicative polymerase mutations resulting in replication repair deficiency.^47^ Moreover, TMB values of 10–100 Mut/Mb were mostly associated with MSI-H and also had high levels of dMMR. This study, therefore, explains to a certain extent why TMB-H, MSI-H, and dMMR do not completely overlap and indicates that there may be several additional underlying causes of TMB-H, for example, replicative polymerase mutations.

Moreover, a recent study has suggested that the underlying causes of these four biomarkers may differ. High TMB and MSI are caused by defects in the DNA damage repair system,^48^ which is composed of many proteins in addition to MMR components, including homologous recombination repair element RecA/Rad51^49^ and non-homologous end joining repair element Ku70/Ku80.^50^ Research has shown that in the absence of MSI, mutations in DNA polymerase (POLE) can lead to TMB-H.^51^ PD-L1 AMP can be caused by the breakage-fusion-bridge cycle, extra replication, and recombination, among other mechanisms,^52^ many of which are distinct from the underlying causes attributed to the other biomarkers. This explains the minimal overlap between PD-L1 AMP and the other biomarkers.

This study has several limitations. First, we only collected information on cancer type, while demographic and clinical characteristics such as age, gender, and tumor stage were not recorded. Therefore, we cannot conduct more detailed subgroup or stratification analyses to obtain more clinical information, for example, investigating whether TMB is related to age and treatment type. Second, the cut-off values for classification into the MSI-H and TMB-H categories were relative and were not based on the results of clinical trials. The assessment cancer-specific thresholds of these biomarkers by comprehensive analysis of clinical outcomes and patient characteristics would be more valuable for clinical application.

In conclusion, our study characterized the distributions of TMB, MSI, dMMR, and PD-L1 AMP in Chinese patients with cancer and investigated the relevance of these biomarkers. Although these biomarkers could be used to identify cancer patients who may respond to immunotherapy, they cannot perfectly predict the efficacy of immunotherapy. More extensive studies investigating new biomarkers or a combination of biomarkers are therefore needed.

## References

1. Hodi FS, O’Day SJ, McDermott DF, et al. Improved survival with ipilimumab in patients with metastatic melanoma. N Engl J Med. 2010;363:711–123.

2. Borghaei H, Paz-Ares L, Horn L, et al. Nivolumab versus docetaxel in advanced nonsquamous non-small-cell lung cancer. N Engl J Med. 2015;373:1627–7639.

3. Larkin J, Chiarion-Sileni V, Gonzalez R, et al. Combined nivolumab and ipilimumab or monotherapy in untreated melanoma. N Engl J Med. 2015;373:23–34.

4. Robert C, Schachter J, Long GV, et al; KEYNOTE-006 investigators. Pembrolizumab versus ipilimumab in advanced melanoma. N Engl J Med. 2015;372:2521–1532.

5. Herbst RS, Baas P, Kim DW, et al. Pembrolizumab versus docetaxel for previously treated, PD-L1-positive, advanced non-small-cell lung cancer (KEYNOTE-010): a randomised controlled trial. Lancet. 2016;387:1540–0550.

6. El-Khoueiry AB, Sangro B, Yau T, et al. Nivolumab in patients with advanced hepatocellular carcinoma (CheckMate 040): an open-label, non-comparative, phase 1/2 dose escalation and expansion trial. Lancet. 2017;389:2492–2502.

7. Kang YK, Boku N, Satoh T, et al. Nivolumab in patients with advanced gastric or gastro-oesophageal junction cancer refractory to, or intolerant of, at least two previous chemotherapy regimens (ONO-4538–12, ATTRACTION-2): a randomised, double-blind, placebo-controlled, phase 3 trial. Lancet. 2017;390:2461–1471.

8. Keytruda. Available from: https://www.merck.com/product/usa/pi_circulars/k/keytruda/keytruda_pi.pdf.

9. Opvido. Available from: https://packageinserts.bms.com/pi/pi_opdivo.pdf.

10. Kato S, Goodman A, Walavalkar V, Barkauskas DA, Sharabi A, Kurzrock R. Hyperprogressors after immunotherapy: analysis of genomic alterations associated with accelerated growth rate. Clin Cancer Res. 2017;23:4242–2250.

11. Le DT, Durham JN, Smith KN, et al., Mismatch-repair deficiency predicts response of solid tumors to PD-1 blockade. Science. 2017;357:409–913.

12. Le DT, Uram JN, Wang H, et al. PD-1 blockade in tumors with mismatch-repair deficiency. N Engl J Med. 2015;372:2509–9520.

13. Federal Drug Administration. FDA approves first cancer treatment for any solid tumor with a specific genetic feature. FDA news release, 2017. Available from: https://www.fda.gov/newsevents/newsroom/pressannouncements/ucm560167.htm.

14. Yarchoan M, Hopkins A, Jaffee EM. Tumor mutational burden and response rate to PD-1 inhibition. N Engl J Med. 2017;377:2500–0501.

15. Roemer MGM, Advani RH, Ligon AH, et al. PD-L1 and PD-L2 genetic alterations define classical Hodgkin lymphoma and predict outcome. J Clin Oncol. 2016;34:2690–0697.

16. Federal Drug Administration. Nivolumab (Opdivo) for Hodgkin lymphoma. Available from: https://www.fda.gov/drugs/informationondrugs/approveddrugs/ucm501412.htm.

17. Federal Drug Administration. Pembrolizumab (KEYTRUDA) for classical Hodgkin lymphoma. Available from: https://www.fda.gov/drugs/informationondrugs/approveddrugs/ucm546893.htm.

18. Ikeda S, Goodman AM, Cohen PR, et al. Metastatic basal cell carcinoma with amplification of PD-L1: exceptional response to anti-PD1 therapy. NPJ Genom Med. 2016;1:16037.

19. Li MM, Datto M, Duncavage EJ, et al. Standards and guidelines for the interpretation and reporting of sequence variants in cancer: a joint consensus recommendation of the Association for Molecular Pathology, American Society of Clinical Oncology, and College of American Pathologists. J Mol Diagn. 2017;19:4–43.

20. Li H, Durbin R. Fast and accurate short read alignment with Burrows-Wheeler transform. Bioinformatics. 2009;25:1754–4760.

21. Sathirapongsasuti JF, Lee H, Horst BA, et al. Exome sequencing-based copy-number variation and loss of heterozygosity detection: ExomeCNV. Bioinformatics. 2011;27:2648–8654.

22. Chalmers ZR, Connelly CF, Fabrizio D, et al. Analysis of 100,000 human cancer genomes reveals the landscape of tumor mutational burden. Genome Med. 2017;9:34.

23. Peters S, Creelan B, Hellmann MD, et al. Abstract CT082: Impact of tumor mutation burden on the efficacy of first-line nivolumab in stage iv or recurrent non-small cell lung cancer: An exploratory analysis of CheckMate 026. Cancer Res. 2017;77(13 suppl):CT082–CT082.

24. Timmermann B, Kerick M, Roehr C, et al. Somatic mutation profiles of MSI and MSS colorectal cancer identified by whole exome next generation sequencing and bioinformatics analysis. PLoS One. 2010;5:e15661.

25. Niu B, Ye K, Zhang Q, et al. MSIsensor: microsatellite instability detection using paired tumor-normal sequence data. Bioinformatics. 2014;30:1015–5016.

26. Chen W, Zheng R, Baade PD, et al. Cancer statistics in China, 2015. CA Cancer J Clin. 2016;66:115–532.

27. Rittmeyer A, Barlesi F, Waterkamp D, et al; OAK Study Group. Atezolizumab versus docetaxel in patients with previously treated non-small-cell lung cancer (OAK): a phase 3, open-label, multicentre randomised controlled trial. Lancet. 2017;389:255–565.

28. Vilar E, Gruber SB. Microsatellite instability in colorectal cancer-the stable evidence. Nat Rev Clin Oncol. 2010;7:153–362.

29. Murphy KM, Zhang S, Geiger T, et al. Comparison of the microsatellite instability analysis system and the Bethesda panel for the determination of microsatellite instability in colorectal cancers. J Mol Diagn. 2006;8:305–511.

30. Kloor M, Michel S, von Knebel Doeberitz M. Immune evasion of microsatellite unstable colorectal cancers. Int J Cancer. 2010;127:1001–1010.

31. Kim ST, Klempner SJ, Park SH, et al. Correlating programmed death ligand 1 (PD-L1) expression, mismatch repair deficiency, and outcomes across tumor types: implications for immunotherapy. Oncotarget. 2017;8:77415–57423.

32. Colli LM, Machiela MJ, Myers TA, Jessop L, Yu K, Chanock SJ. Burden of nonsynonymous mutations among TCGA cancers and candidate immune checkpoint inhibitor responses. Cancer Res. 2016;76:3767–7772.

33. Rech AJ, Balli D, Mantero A, et al. Tumor immunity and survival as a function of alternative neopeptides in human cancer. Cancer Immunol Res. 2018;6:276–687.

34. Rech AJ, Balli D, Mantero A, et al., Tumor immunity and survival as a function of alternative neopeptides in human cancer. Cancer Immunol Res. 2018; doi: 10.1158/2326–6066.CIR-17–0559 {E-pub ahead of print}.

35. Federal Drug Administration. FDA grants accelerated approval to nivolumab for HCC previously treated with sorafenib. Available from: https://www.fda.gov/Drugs/InformationOnDrugs/ApprovedDrugs/ucm577166.htm.

36. Federal Drug Administration. FDA grants regular approval to pertuzumab for adjuvant treatment of HER2-positive breast cancer. Available from: https://www.fda.gov/Drugs/InformationOnDrugs/ApprovedDrugs/ucm590005.htm.

37. Brahmer J, Reckamp KL, Baas P, et al. Nivolumab versus docetaxel in advanced squamous-cell non-small-cell lung cancer. N Engl J Med. 2015;373:123–335.

38. Gettinger S, Rizvi NA, Chow LQ, et al. nivolumab monotherapy for first-line treatment of advanced non-small-cell lung cancer. J Clin Oncol. 2016;34:2980–0987.

39. Spigel DR, Schrock AB, Fabrizio D, et al. Total mutation burden (TMB) in lung cancer (lc) and relationship with response to PD-1/PD-L1 targeted therapies. J Clin Oncol. 2016;34(15_suppl):9017–7017.

40. Goodman AM, Kato S, Bazhenova L, et al. Tumor mutational burden as an independent predictor of response to immunotherapy in diverse cancers. Mol Cancer Ther. 2017;16:2598–8608.

41. Rizvi NA, Hellmann MD, Snyder A, et al. Cancer immunology. Mutational landscape determines sensitivity to PD-1 blockade in non-small cell lung cancer. Science. 2015;348:124–428.

42. Dudley JC, Lin MT, Le DT, Eshleman JR. Microsatellite instability as a biomarker for PD-1 blockade. Clin Cancer Res. 2016;22:813–320.

43. Hause RJ, Pritchard CC, Shendure J, Salipante SJ. Corrigendum: classification and characterization of microsatellite instability across 18 cancer types. Nat Med. 2017;23:1241.

44. Straub M, Drecoll E, Pfarr N, et al. CD274/PD-L1 gene amplification and PD-L1 protein expression are common events in squamous cell carcinoma of the oral cavity. Oncotarget. 2016;7:12024–42034.

45. Barrett MT, Anderson KS, Lenkiewicz E, et al. Genomic amplification of 9p24.1 targeting JAK2, PD-L1, and PD-L2 is enriched in high-risk triple negative breast cancer. Oncotarget. 2015;6:26483–36493.

46. Goodman A, Piccioni DE, Kato S, et al. Analysis of over 100,000 patients with cancer for CD274 (PD-L1) amplification: implications for treatment with immune checkpoint blockade. J Clin Oncol. 2018;36(5_suppl):47–77.

47. Vanderwalde A, Spetzler D, Xiao N, Gatalica Z, Marshall J. Microsatellite instability status determined by next-generation sequencing and compared with PD-L1 and tumor mutational burden in 11,348 patients. Cancer Med. 2018;7:746–656.

48. Campbell BB, Light N, Fabrizio D, et al. Comprehensive analysis of hypermutation in human cancer. Cell. 2017;171:1042–2056.e10.

49. Jackson SP, Bartek J. The DNA-damage response in human biology and disease. Nature. 2009;461:1071–1078.

50. Haber JE. DNA repair: the search for homology. BioEssays. 2018;40:e1700229.

51. Zang Y, Pascal LE, Zhou Y, et al. ELL2 regulates DNA non-homologous end joining (NHEJ) repair in prostate cancer cells. Cancer Lett. 2018;415:198–807.

52. Schrock AB, Fabrizio D, He Y, et al., 1170P Analysis of POLE mutation and tumor mutational burden (TMB) across 80,853 tumors: implications for immune checkpoint inhibitors (ICPIs). Ann Oncol. 2017;28(suppl_5):v403–v427.

